# Restored rigidity sensing fails to induce anoikis in cancer cells due to Akt activation and cell–cell interactions

**DOI:** 10.1101/2025.06.11.659173

**Authors:** Anat Galis Vivante, Nehal Dwivedi, Michael P. Sheetz, Guy Nir

## Abstract

Rigidity sensing enables cells to detect and respond to extracellular matrix stiffness, directing survival and apoptotic decisions. Tropomyosin 2.1 (Tpm2.1), a key actin-binding protein, is essential for this process and is frequently downregulated in cancer. While Tpm2.1 overexpression restores rigidity sensing and suppresses malignant growth, the downstream pathways governing cell fate remain poorly defined. Here, we show that Tpm2.1-mediated rigidity sensing represses anchorage-independent growth by downregulating oncogenic networks, including PI3K–Akt and EphA2 signaling. Unexpectedly, a majority of Tpm2.1-overexpressing cells remained viable in suspension, revealing a previously unrecognized decoupling between rigidity sensing and anoikis. Transcriptomic profiling of apoptotic and non-apoptotic cells in suspension identified compensatory survival programs marked by PI3K–Akt activation and induction of cell–cell adhesion genes. These findings uncover a dominant anoikis-resistant state that persists despite cytoskeletal normalization, suggesting that rigidity sensing alone is insufficient to reestablish anchorage dependence in cancer cells.

## Main

Anchorage-independent growth is a hallmark of metastatic cancer, allowing tumor cells to survive and proliferate without attachment to the extracellular matrix (ECM)^1–4^. This capacity is closely linked to defects in rigidity sensing, a cellular process in which anchorage-dependent cells interpret ECM stiffness to regulate survival, proliferation, and apoptosis^5–7^. In normal tissues, detachment from the ECM activates a specialized form of programmed cell death known as anoikis, thereby preventing inappropriate growth in suspension^8–11^. In contrast, metastatic cells evade anoikis, enabling dissemination through circulation and colonization of distant sites^12^.

Rigidity sensing is mediated through focal adhesions, which are macromolecular complexes that connect integrin clusters at the membrane to the actomyosin cytoskeleton^13–16^. On rigid ECM, elevated contractility stabilizes focal adhesions, promoting cytoskeletal organization and pro-survival signaling. Conversely, on soft ECM, low mechanical resistance prevents adhesion maturation and activates apoptotic pathways^17–21^. The actin-binding protein Tropomyosin 2.1 (Tpm2.1) plays a key role in this process. Frequently downregulated in cancer^22, 23^, Tpm2.1 is essential for proper rigidity sensing and cytoskeletal integrity. Its re-expression in malignant cells has been shown to restore rigidity sensing, suppress anchorage-independent growth, and induce apoptosis when cells are cultured in suspension^6, 11, 24^.

Although integrin-ECM signaling activates survival pathways such as focal adhesion kinase (FAK) and MAPK^25–27^, it remains unclear whether restoring rigidity is sufficient to overcome oncogenic adaptations that promote survival in suspension. Cancer cells adapted to anchorage independence exhibit widespread transcriptional changes, including suppression of integrin–ECM signaling, inhibition of the Hippo-YAP/TEAD axis, and increased expression of anti-apoptotic mediators such as clusterin and TNF-related factors^28–30^. These reprogramming events are thought to confer resistance to anoikis in the absence of mechanical cues^31, 32^. While re-expression of Tpm2.1 restores rigidity sensing and can induce apoptosis on soft matrices or in suspension^33–35^, whether this is sufficient to eliminate anchorage-independent survival has not been systematically evaluated in dynamic or heterogeneous cell populations. Here, we demonstrate that despite restored rigidity sensing, the majority of cells remain viable in suspension, revealing alternative transcriptional programs that maintain survival independently of matrix attachment.

To uncover the molecular basis of anoikis resistance in cells with restored rigidity sensing, we used MDA-MB-231 triple-negative breast cancer (TNBC) cells, which endogenously express low levels of Tpm2.1. We compared them to the same line engineered to overexpress Tpm2.1^35^. Using time-resolved transcriptomic profiling, we characterized the transition from adherent to suspension states under conditions of impaired or restored rigidity sensing. Surprisingly, most Tpm2.1-overexpressing cells persisted in suspension, revealing a decoupling between rigidity sensing and anoikis susceptibility. Mechanistically, these cells escaped early apoptosis by activating NF-κB signaling and inducing cell adhesion molecules (CAMs) that promote cell–cell interactions^36^. By day four, this compensatory program converged on activation of the PI3K–Akt pathway, a key mediator of TNBC survival and therapeutic resistance^37–40^. Together, these findings identify an unanticipated, dominant anoikis-resistant state in mechanically responsive cells, and suggest new therapeutic vulnerabilities in metastatic progression.

## Results

### Tpm2.1 expression promotes anoikis and inhibits migration in anoikis-resistant MDA-MB-231 breast cancer cells

MDA-MB-231 cells express low levels of Tpm2.1, a deficiency linked to their ability to survive and proliferate on soft matrices^6, 11^. To assess whether Tpm2.1 re-expression promotes apoptosis in suspension, we cultured wild-type MDA-MB-231 cells (MDA-WT), cells stably expressing an empty vector (MDA-EV), and cells stably expressing Tpm2.1 (MDA-Tpm2.1)^35^ on ultra-low attachment (ULA) plates (see Methods), which prevent substrate adhesion and force cells into suspension^41^. Growth in non-adherent conditions is a hallmark of cellular transformation^1^ and correlates strongly with outcomes in soft agar assays^42^. To capture dynamic changes following detachment, we conducted a time-course experiment in which cells were maintained either in attached or in suspension conditions for 5 hours, 1 day, 2 days, or 4 days (Fig. 1A).

**Figure 1.**
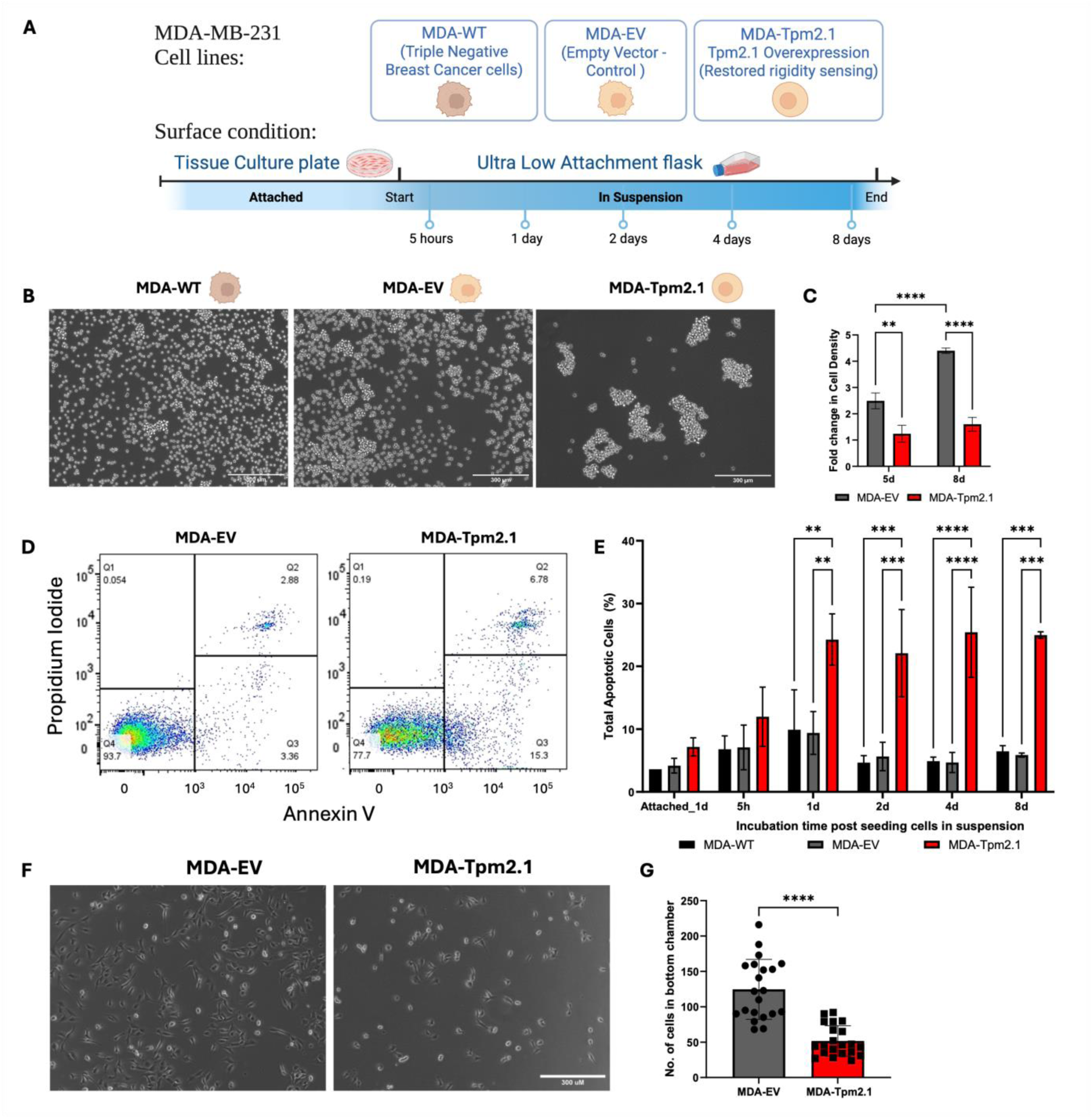
Tpm2.1 expression promotes anoikis in anoikis-resistant MDA-MB-231 breast cancer cells. (A) Schematic of MDA-EV and MDA-Tpm2.1 cells cultured under adherent (tissue culture plate) and suspension (low-attachment plate) conditions. (B) Representative phase-contrast images of MDA-WT, MDA-EV and MDA-Tpm2.1 cells after 8 days in suspension culture. (C) Mean ± SD of fold change in cell density for MDA-Tpm2.1 and MDA-EV cells cultured in suspension for 5 and 8 days. (D) Pseudocolor flow cytometry plots of Annexin V and propidium iodide (PI) staining in cells cultured in suspension for 4 days show increased apoptosis in MDA-Tpm2.1 cells. Quadrants reflect populations enriched for live, early apoptotic, and late apoptotic or dead cells based on Annexin V and PI signal intensities. (E) Mean ± SD of total apoptotic cells (%) in attached (1 day) and suspension at 5 h, 1 day, 2 days, 4 days, and 8 days. (F) Representative phase images of MDA-EV and MDA-Tpm2.1 cells migrated to the bottom well after 48 h in the trans-well chamber. (G) Mean ± SD of the total number of migrated cells. (**) for p value < 0.01, (***) for p value < 0.001, (****) for p value <0.0001.

**Figure S1.**
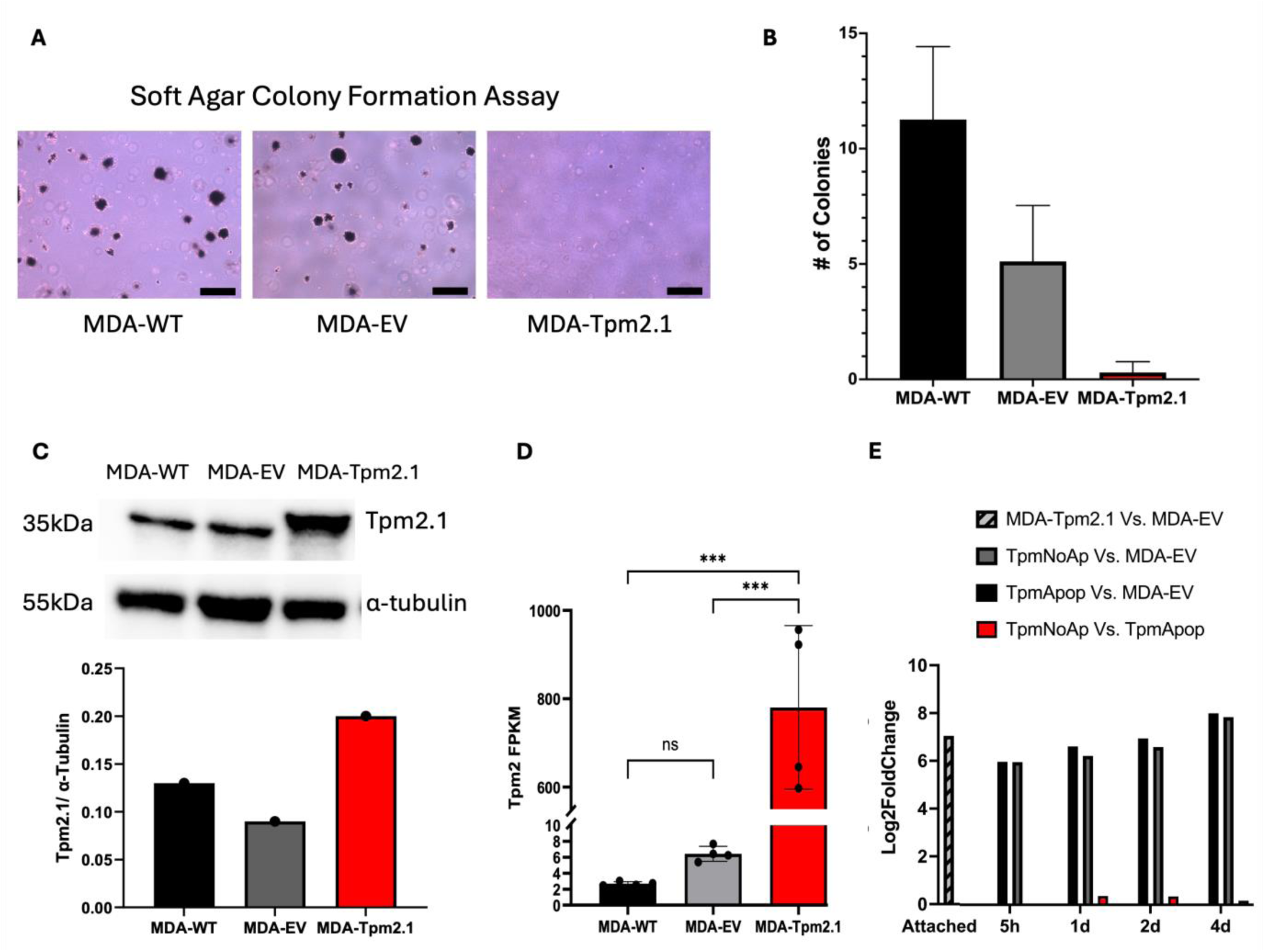
Tpm2.1 expression enhances anoikis in MDA-MB-231 breast cancer cells - Control experiments. (A) Soft agar colony formation assay to ensure that MDA-Tpm2.1 cells do not grow on soft surfaces. Representative phase-contrast images of MDA-WT, MDA-EV, and MDA-Tpm2.1 colonies in 3D soft agar. Scale bar: 100 μm. (B) Quantification (mean ± SD) of colony number for MDA-WT, MDA-EV, and MDA-Tpm2.1 cells grown in soft agar. Colonies were stained with crystal violet, counted, and averaged from three biological replicates (n = 3). (C) Western blot analysis and quantification of Tpm2.1 protein expression in MDA-WT, MDA-EV, and MDA-Tpm2.1 cells. (D) Bar graph (mean ± SD) showing fragments per kilobase of transcript per million mapped reads (FPKM) for Tpm2 expression in MDA-WT, MDA-EV, and MDA-Tpm2.1 cells cultured under attached conditions, as determined by RNA sequencing. (E) RNA-seq analysis of Tpm2.1 gene expression in MDA-EV and MDA-Tpm2.1 cells following apoptotic selection at various time points during suspension culture.

We monitored cell proliferation and apoptosis during suspension culture (Fig. 1B–E) and assessed colony formation in soft agar (Fig. S1A, B). Both MDA-WT and MDA-EV cells survived and proliferated under suspension conditions and in soft agar, demonstrating anchorage-independent growth. In contrast, MDA-Tpm2.1 cells did not survive in soft agar, indicating anchorage-dependent growth. MDA-WT and MDA-EV cells formed significantly more colonies than MDA-Tpm2.1 cells (Fig. S1A, B). MDA-EV cells exhibited robust growth in suspension, with a 2.5-fold increase in cell number by day 5 and a 4.4-fold increase by day 8. In contrast, MDA-Tpm2.1 cells showed markedly reduced proliferation (1.2-fold and 1.6-fold at days 5 and 8, respectively) (Fig. 1B, C). Apoptosis assays revealed that Tpm2.1 expression significantly increased anoikis compared to both MDA-EV and MDA-WT cells, with elevated apoptosis detectable as early as day 1 and peaking at day 4 (Fig. 1D, E). Notably, despite this increase, only ∼30% of MDA-Tpm2.1 cells were apoptotic by day 4, indicating that approximately 70% of cells remained viable under suspension conditions. RNA-seq and western blot data show that Tpm2.1 expression levels are comparable in both MDA-WT and MDA-EV cells (Fig. S1C, D). Additionally, the levels of apoptosis under suspension conditions were similar between MDA-WT and MDA-EV cells (Fig 1E). Therefore, we used MDA-EV cells as a control for all subsequent experiments.

Given that MDA-MB-231 is a highly metastatic cell line^43, 44^, we next asked whether Tpm2.1 expression also impairs migratory behavior. Transwell migration assays^45^ showed that MDA-Tpm2.1 cells exhibited significantly reduced motility compared to MDA-EV controls (Fig. 1F, G).

Together, these results demonstrate that Tpm2.1 re-expression partially restores anoikis sensitivity and suppresses anchorage-independent growth and migration in metastatic breast cancer cells. However, the persistence of a large non-apoptotic population in suspension suggests the presence of compensatory survival mechanisms that override mechanical cues.

### Transcriptomic profiling reveals distinct gene expression programs in apoptotic and non-apoptotic Tpm2.1-expressing cells

Tpm2.1 expression has been identified as a key regulator of mechanosensing in MDA-MB-231 cells. However, as shown above, despite Tpm2.1 overexpression, a substantial fraction of cells remains viable under non-adherent conditions, indicating incomplete restoration of anoikis sensitivity. To investigate how Tpm2.1 influences the transition to distinct physiological states, we examined the gene expression programs associated with apoptotic versus non-apoptotic responses during suspension culture.

At each time point, MDA-Tpm2.1 cells were separated into two subpopulations: Annexin V-negative (non-apoptotic, Tpm_NoAp) and Annexin V-positive (apoptotic, Tpm_Apop) cells. Cells were sorted using magnetic separation with an Annexin V Microbead Kit^46^, which isolates apoptotic cells by binding Annexin V-labeled microbeads followed by magnetic column selection (Fig. 2A).

**Figure 2.**
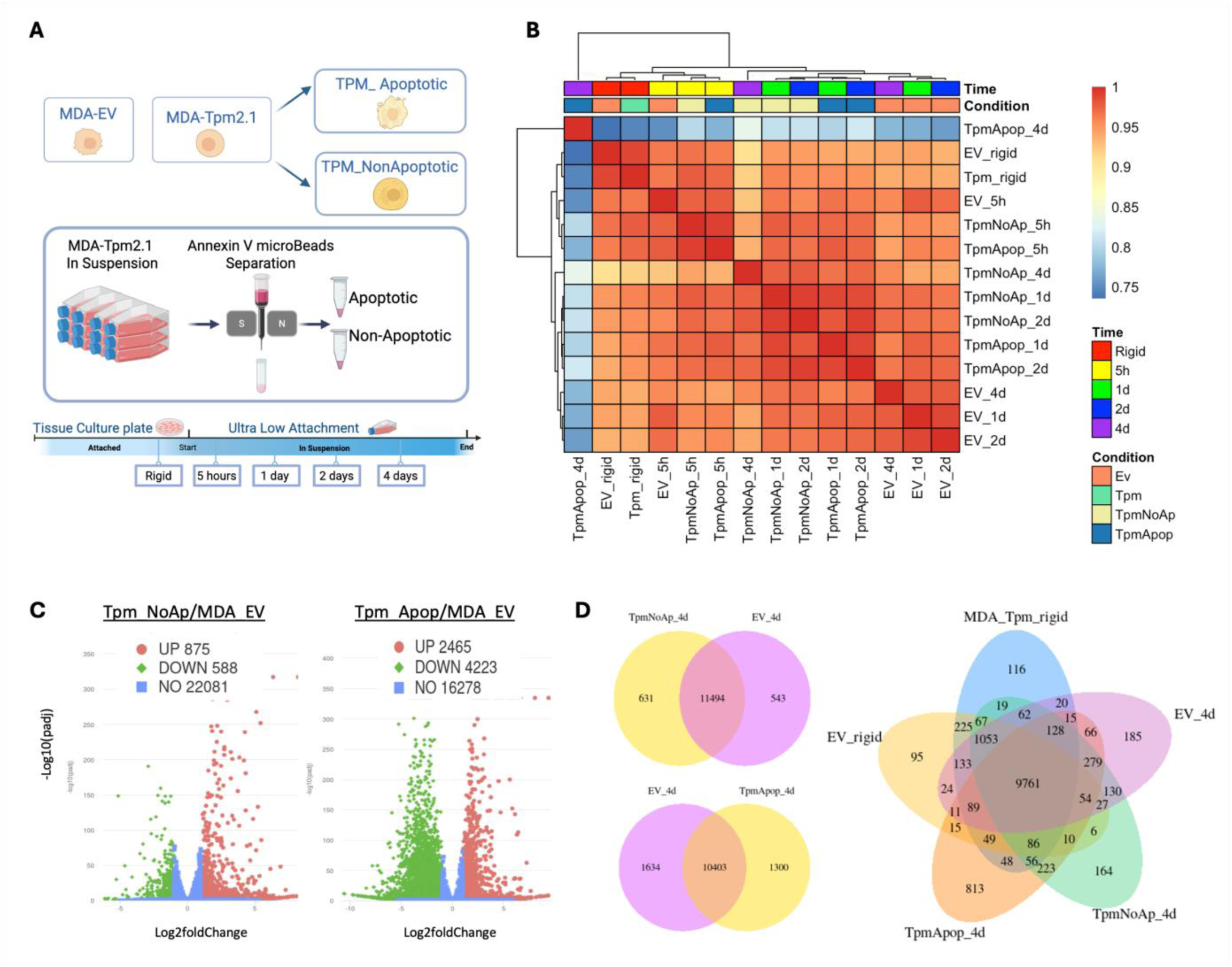
Transcriptomic profiling of apoptotic and non-apoptotic subpopulations in Tpm2.1-expressing MDA-MB-231 cells under suspension conditions. (A) Schematic of the experimental design. MDA-MB-231 cells stably expressing an empty vector (MDA-EV) or Tpm2.1 (MDA-Tpm2.1) were cultured under adherent or suspension conditions. At each suspension time point (5 hours, 1 day, 2 days, and 4 days), MDA-Tpm2.1 cells were separated into apoptotic (Annexin V-positive) and non-apoptotic (Annexin V-negative) subpopulations using Annexin V-conjugated magnetic microbeads, followed by transcriptomic profiling. (B) Spearman correlation heatmap of all RNA-seq samples across time points and conditions. Correlation strength is color-coded, with red indicating strong positive correlation and blue indicating strong negative correlation. The transcriptional profile of apoptotic Tpm2.1-expressing cells at day 4 (Tpm_Apop_4d) is most distinct from all other conditions. (C) Volcano plots showing differentially expressed genes between Tpm_NoAp and Tpm_Apop populations relative to MDA-EV at day 4. Red and green dots represent significantly upregulated and downregulated genes, respectively (adjusted p < 0.05, |log₂FoldChange| > 1). (D) Venn diagram showing the number and overlap of differentially expressed genes across multiple comparisons: Tpm_NoAp vs. MDA-EV, Tpm_Apop vs. MDA-EV, MDA-Tpm2.1 (bulk) vs. MDA-EV, and MDA-EV under rigid conditions vs. day 4 suspension. Differential expression analysis was performed using Novogene’s Novomagic platform, employing the DESeq2 pipeline [adjusted p-value (padj) < 0.05 and |log₂FoldChange| > 1] as significance thresholds.

We then performed time-resolved RNA sequencing of MDA-EV, MDA-Tpm2.1, Tpm_NoAp, and Tpm_Apop cells. Transcriptomic analysis revealed widespread gene expression changes over time and across conditions. A Spearman correlation heatmap (Fig. 2B) showed that the most distinct transcriptional divergence emerged at day 4, when Tpm_Apop cells displayed a gene expression profile markedly different from all other samples. Early time points (rigid and 5-hour suspension) showed minimal differences between MDA-EV and MDA-Tpm2.1 cells, suggesting that short-term detachment does not yet invoke a Tpm2.1-specific response. In contrast, longer suspension periods (≥1 day) revealed progressive separation of Tpm2.1-expressing cells from controls, with the strongest effect observed in the apoptotic subpopulation at day 4.

To confirm that these differences were not due to variation in Tpm2.1 expression itself, we measured Tpm2.1 transcript levels across all conditions and found them to be stable (Fig. S1E). Differential gene expression analysis, performed using the Novomagic platform (see Methods), revealed thousands of significantly altered genes when comparing Tpm_Apop to MDA-EV cells. By contrast, the Tpm_NoAp population showed only a few hundred differentially expressed genes relative to MDA-EV (Fig. 2C, D), suggesting that the apoptotic state in Tpm2.1-overexpressing cells is accompanied by a dramatic and distinct transcriptional shift.

### Distinct transcriptional programs underlie survival and death in suspended MDA-MB-231 cells

To identify the signaling pathways associated with anoikis resistance or susceptibility, we performed Kyoto Encyclopedia of Genes and Genomes (KEGG)^47^ pathway enrichment analysis across all time points (5 h, 1 d, 2 d, and 4 d) in MDA-EV, Tpm_NoAp, and Tpm_Apop. Despite all groups experiencing suspension stress, the three cell populations activated markedly different transcriptional programs that corresponded to their fate (Fig. 3A and Supplementary Information).

**Figure 3.**
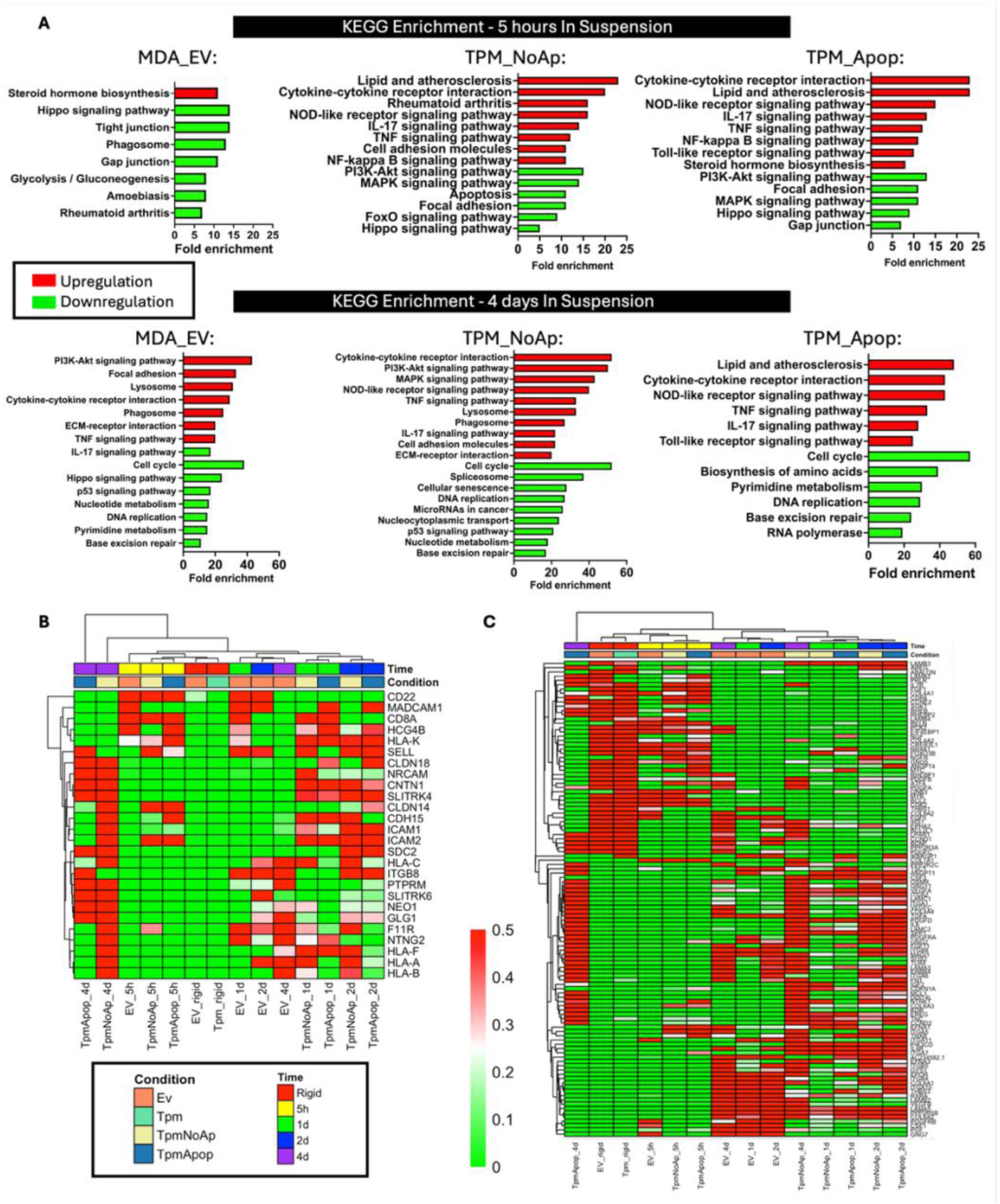
KEGG pathway enrichment analyses for MDA-MB-231 Cancer Cells Induced by Tpm2.1 Overexpression and Suspension Growth Conditions. (A) KEGG pathway enrichment analysis of upregulated and downregulated genes in MDA-EV, Tpm_NoAp, and Tpm_Apop cells after 5 hours and 4 days in suspension compared to their respective cell populations cultured in attached (rigid) conditions. Red bars represent upregulated genes. Green bars represent downregulated genes. All pathways have adjusted p < 0.05, |log₂FoldChange| > 1. (B, C) Heatmaps of differentially expressed genes from KEGG analysis for the Cell Adhesion Molecules (CAM) pathway (B) and the PI3K-Akt signaling pathway (C). Expression levels are color-coded, with red indicating high expression and green indicating low expression. Heatmaps were generated using an R script, with adjusted p-value (padj) < 0.05 and |log₂FoldChange| > 1, applying Spearman correlation and the centroid method for clustering.

MDA-EV cells, which proliferate in suspension, rapidly adapted to detachment by upregulating steroid hormone biosynthesis, likely reflecting lipid-based compensatory signaling^48, 49^. This early adaptation was accompanied by suppression of cell–cell communication pathways (e.g., Hippo, gap junctions, tight junctions), suggestive of disengagement from multicellular coordination^50–52^. Over time, MDA-EV cells progressively reactivated ECM–receptor interaction^53^, focal adhesion^54^, and the PI3K–Akt^55^ signaling pathway, supporting anchorage-independent growth^30, 56^. Notably, this occurred in parallel with downregulation of p53, DNA replication, and cell cycle pathways, indicating a decoupling from canonical adhesion-mediated checkpoint control^57^.

Tpm2.1-overexpressing cells that resisted anoikis (Tpm_NoAp) followed a markedly different trajectory. While they also suppressed cell cycle, p53, and Hippo pathways early on, they uniquely mounted a robust inflammatory and immune-related transcriptional response. These include upregulation of TNF, IL-17, NF-κB, and Toll-like receptor (TLR) signaling as early as 5 hours post-detachment. Concomitantly, both Tpm_NoAp and Tpm_Apop cells showed early suppression of the MAPK pathway, known to mediate apoptosis^58, 59^. However, MAPK signaling was selectively reactivated in Tpm_NoAp cells by day 4, potentially contributing to longer-term survival.

A defining feature of the Tpm_NoAp population was the persistent upregulation of the cell adhesion molecules (CAMs) pathway, including genes such as ICAM1, ICAM2, CDH15, F11R, HLA-A, and HLA-B. These promote cell–cell adhesion and immune-related signaling and may collectively support multicellular clustering, enhancing resistance to anoikis^60^. This was further complemented by delayed activation of the PI3K–Akt pathway at 4 days, including elevated expression of ITGA5, ITGA11, ITGB4, CDKN1A, NTRK1, and LAMB3. Genes which mediate adhesion, survival, and tumor progression^61–63^. Together, these data suggest that anoikis resistance in Tpm_NoAp cells is mediated by an integrated transcriptional program that includes early immune priming, reinforced cell–cell connectivity, and late pro-survival signaling.

In contrast, Tpm_Apop cells failed to sustain these compensatory responses. Although they transiently activated inflammatory and steroid biosynthesis pathways, they did not reactivate PI3K–Akt or MAPK signaling at later stages. Instead, they showed persistent downregulation of focal adhesion, cell cycle, DNA repair, and RNA processing pathways. Notably, only Tpm_Apop cells upregulated the TLR pathway at day 4, potentially indicating activation of innate immune-like stress signals linked to cell death^64, 65^. These features suggest a transcriptional trajectory consistent with irreversible commitment to apoptosis.

To further investigate the differential regulation of pro-survival circuits, we focused on the CAM and PI3K–Akt pathways, which emerged as central to the divergence between Tpm_NoAp and Tpm_Apop cells.

Analysis of the CAM pathway (Fig. 3B) revealed elevated expression of intercellular adhesion genes, including F11R, HLA-A, HLA-B, ICAM1, ICAM2, and CDH15, specifically in the Tpm_NoAp population by day 4. These genes support immune signaling and promote tight cell-to-cell adhesion^66, 67^. Their selective upregulation in non-apoptotic cells, but not in the apoptotic population, suggests that collective cohesion may act as a compensatory survival mechanism in the absence of extracellular matrix attachment.

In parallel, expression profiling of the PI3K–Akt pathway (Fig. 3C) showed upregulation of integrin subunits (ITGA5, ITGA11, ITGB4) and additional adhesion and survival-related genes, including TNXB, CSF3, CDKN1A, NTRK1, and LAMB3, in MDA-EV and Tpm_NoAp cells. These genes were not enriched in the apoptotic group, reinforcing the idea that the PI3K–Akt axis is selectively activated in populations capable of long-term survival in suspension.

Together, these pathway-level analyses underscore how non-apoptotic cells engage both adhesive and signaling-based mechanisms to maintain viability after detachment, while apoptotic cells fail to activate these compensatory programs.

### Cell–cell interactions contribute to anoikis resistance in Tpm2.1-expressing cells

The pathway enrichment analysis led us to hypothesize that upregulation of cell–cell interactions may promote survival in Tpm_NoAp cells during anchorage-independent growth.

We therefore seeded cells at low (1 × 10³ cells/well), medium (5 × 10^4^ cells/well), and high (1 × 10⁶ cells/well) densities in low-attachment plates and measured apoptosis after four days in suspension (Fig. 4A, B). MDA-Tpm2.1 cells exhibited significantly higher apoptosis levels at low cell density, which was markedly reduced at higher densities. This indicates that cell-cell communication supports survival. In contrast, MDA-EV cells showed no appreciable change in apoptosis across different seeding densities.

**Figure 4.**
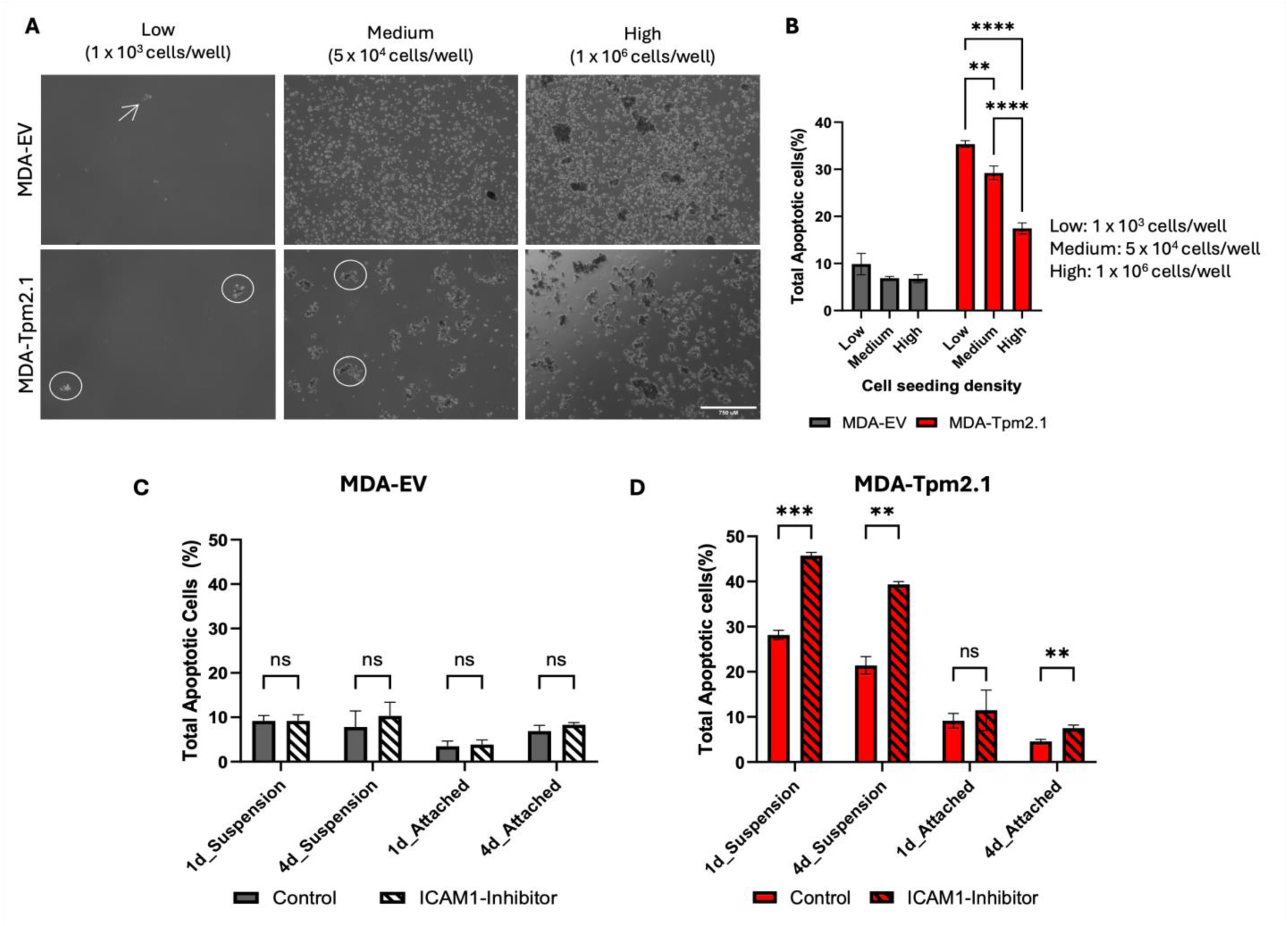
Change in cell-cell interaction affects anoikis in MDA-Tpm2.1 cells. (A) Phase-contrast images of MDA-EV and MDA-Tpm2.1 cells cultured at three different cell densities for 4 days in suspension. MDA-EV cells are indicated by arrows; MDA-Tpm2.1 cell aggregates are highlighted by circles. (B) Mean ± SD of total apoptotic cells (Annexin V^+^ only, PI^+^ only, and Annexin V^+^/PI^+^; %) in suspension cultures at different cell densities for 4 days. (C) Phase-contrast images of MDA-Tpm2.1 cells treated with DMSO (control) or ICAM-1 inhibitor (100nM) for 1 day and 4 days in suspension culture. (D) Mean ± SD of total apoptotic cells (Annexin V^+^ only, PI^+^ only, and Annexin V^+^/PI^+^; %) in MDA-EV (grey) and MDA-Tpm2.1(red) cells treated with ICAM-1 inhibitor (ICAM-1-IN-1, 100nM) for 1 days or 4 days under suspension and attached conditions. The ICAM-1 inhibitor was added at the time of seeding. (**) for p value < 0.01, (***) for p value < 0.001, (****) for p value <0.0001

Among the upregulated cell adhesion molecules, ICAM-1 showed the highest fold change in non-apoptotic MDA-Tpm2.1 cells under suspension conditions compared to adherent conditions (Fig. 3B). ICAM-1 is known to be overexpressed in circulating tumor cells and has been implicated in promoting breast cancer metastasis^68^. We therefore investigated whether ICAM-1 contributes functionally to the survival of MDA-Tpm2.1 cells in suspension. Inhibition of ICAM-1 led to increased apoptosis in MDA-Tpm2.1 cells at both one and four days in suspension. Under adherent conditions, ICAM-1 inhibition had no significant effect at day 1, though a modest increase in apoptosis was observed by day 4 (Fig. 4C, D).

Collectively, these findings suggest that enhanced cell–cell adhesion contributes to anoikis resistance in Tpm2.1-expressing cells, and that disrupting this interaction, particularly via ICAM-1 inhibition, can sensitize these cells to undergo apoptosis under anchorage-independent conditions.

### Tpm2.1-mediated suppression of AKT and EphA2 signaling promotes anoikis in breast cancer cells

A second key distinction identified in our pathway enrichment analysis was the selective upregulation of the PI3K–AKT signaling pathway in MDA-EV and Tpm_NoAp cells, but not in Tpm_Apop cells, after four days in suspension. Activation of this pathway occurs through phosphorylation of AKT at Threonine 308 by PDK1 and at Serine 473 by mTOR, both of which are essential for full AKT activation^39, 69^. Therefore, we sought to compare the levels of phosphorylated AKT and total AKT in MDA-EV and MDA-Tpm2.1 cells, under both suspension and adherent conditions.

Western blot analysis revealed that phospho-AKT (Thr308) levels were reduced in MDA-Tpm2.1 cells compared to MDA-EV cells during suspension culture, while no such difference was observed under adherent conditions. Total AKT expression decreased in both cell types in suspension relative to adhesion, but levels were comparable between MDA-EV and MDA-Tpm2.1 cells under each condition. These results indicate that Tpm2.1 suppresses AKT activation specifically in suspension, rather than altering its basal expression (Fig. 5A–C).

**Figure 5.**
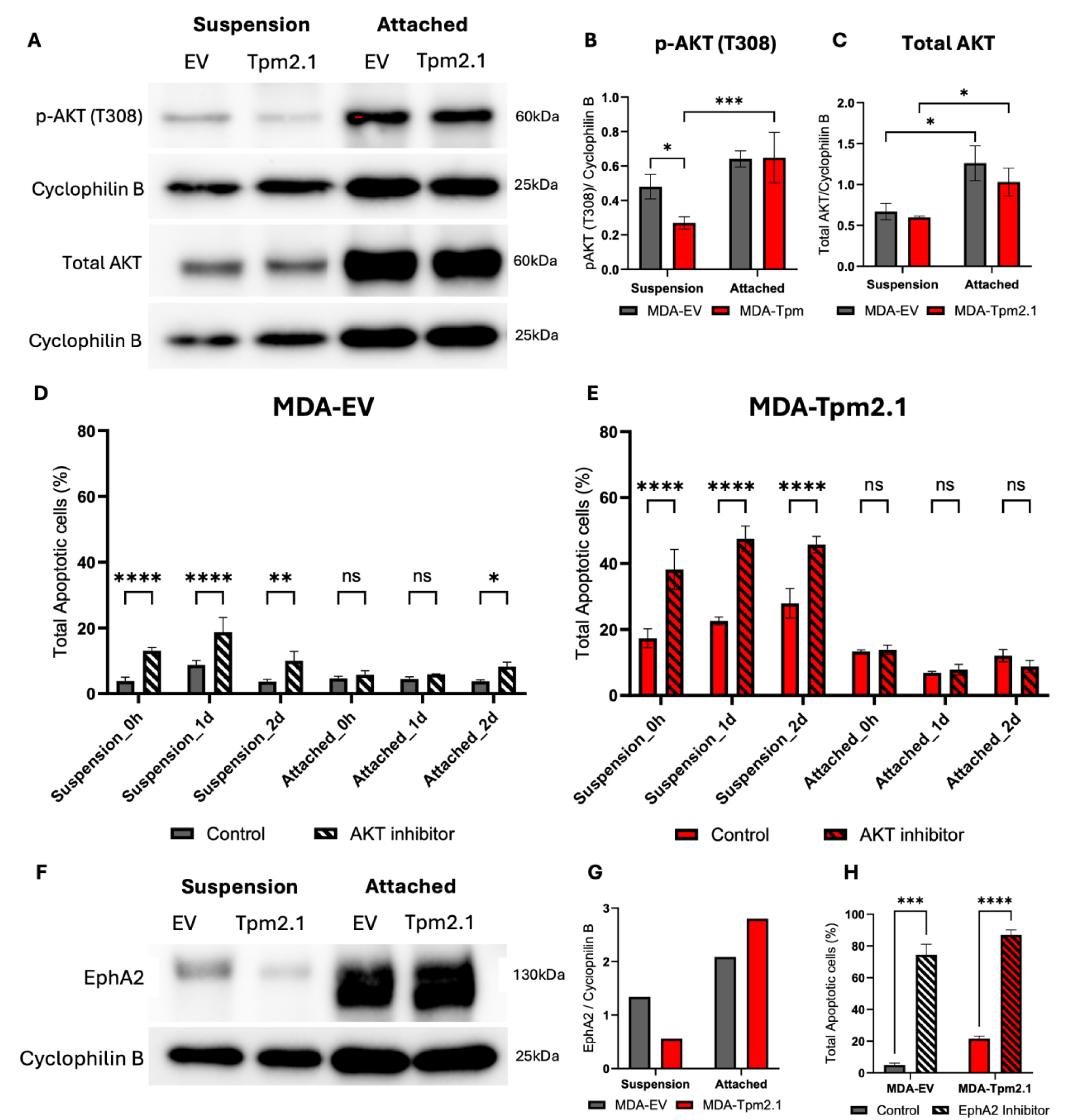
Tpm2.1 expression downregulates the AKT pathway in MDA cells growing in suspension. (A) Western blot showing the expression of phospho-AKT (T308) and total AKT in MDA-EV and MDA-Tpm2.1 cells cultured under suspension and attached conditions for 5 days. (B, C) Quantification of (B) phospho-AKT (T308) and (C) total AKT levels in MDA-EV and MDA-Tpm2.1 cells under suspension and attached conditions for 5 days, presented as mean ± SD. (D, E) Quantification of total apoptotic cells (%) presented as mean ± SD for (D) MDA-EV and (E) MDA-Tpm2.1 cells after treatment with AKT inhibitor (5 µM) for 4 days under suspension and attached conditions. The AKT inhibitor was added at the time of seeding (0 h), 1 day, and 2 days after seeding, as indicated on the X-axis of panels D and E. (F) Western blot showing the expression of EphA2 in MDA-EV and MDA-Tpm2.1 cells cultured under suspension and attached conditions for 5 days. (G) Quantification of EphA2 levels in MDA-EV and MDA-Tpm2.1 cells under suspension and attached conditions for 5 days. N=1. (H) Quantification of total apoptotic cells (%) presented as mean ± SD for MDA-EV and MDA-Tpm2.1 cells after treatment with EphA2 kinase inhibitor (1.5nM) for 4 days under suspension and attached conditions. (*) for p value <0.05, (**) for p value < 0.01, (***) for p value < 0.001, (****) for p value <0.0001.

**Figure S2:**
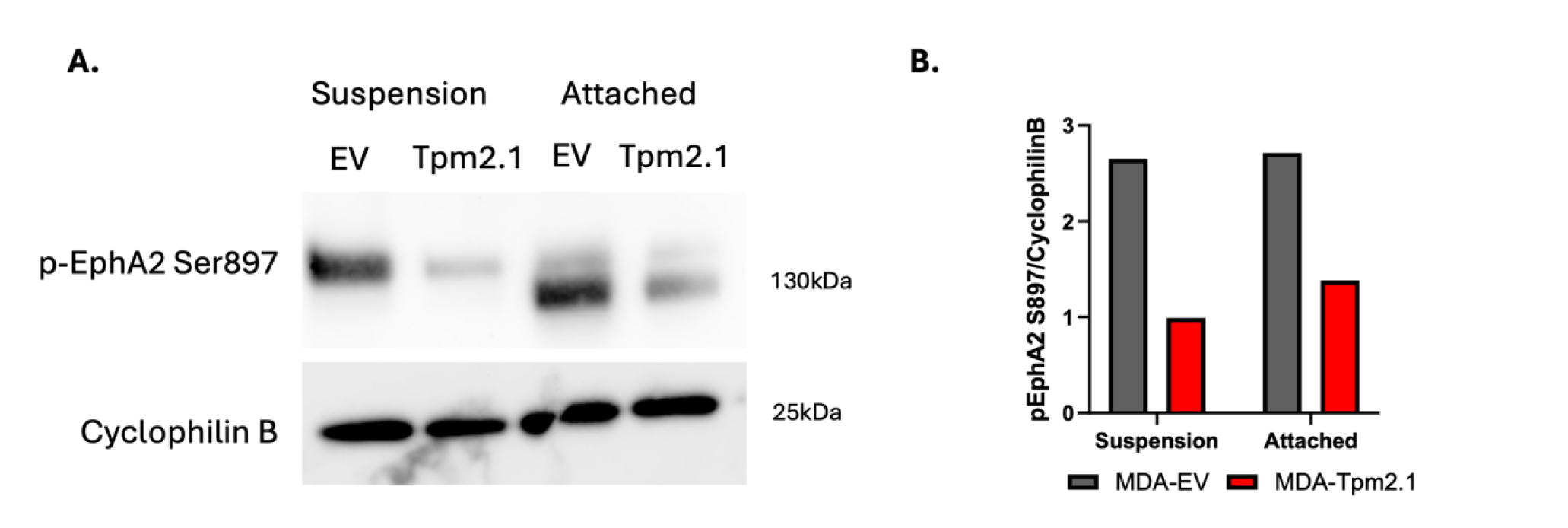
Phospho-EpHA2 (Ser897) levels in MDA-EV and MDA-Tpm2.1 cells. (A) Western blot showing the expression of phospho-EphA2(Ser897) in MDA-EV and MDA-Tpm2.1 cells cultured under suspension and attached conditions for 5 days. (B) Quantification of band intensity for phospho-EphA2(Ser897), normalized to cyclophilin B (loading control) in MDA-EV and MDA-Tpm2.1 cells under suspension and attached conditions for 5 days.

To assess the functional role of AKT signaling in anoikis resistance, we treated the cells with Afuresertib Hydrochloride, an AKT kinase inhibitor, for four days. Since our pathway enrichment analysis revealed that this pathway was upregulated after four days, we sought to determine whether the timing of AKT inhibition influences apoptotic outcomes. Therefore, the inhibitor was added at three time-points: during cell seeding (0 h), one day, and two days post-seeding. Both MDA-EV and MDA-Tpm2.1 cells exhibited increased apoptosis upon AKT inhibition. Notably, MDA-EV cells showed a more pronounced apoptotic response when the inhibitor was added at the time of seeding (Fig. 5D). In contrast, MDA-Tpm2.1 cells displayed similar levels of apoptosis regardless of the timing of inhibitor administration (Fig. 5E).

To further understand the mechanisms by which the AKT pathway supports anoikis resistance, we examined its downstream effectors. Heatmap analysis of the PI3K-AKT pathway revealed elevated expression of EphA2 in MDA-EV cells and in the non-apoptotic population of MDA-Tpm2.1 cells cultured in suspension for four days, compared to the apoptotic fraction of MDA-Tpm2.1 cells (Fig. 3C). Based on this observation, we examined EphA2 protein levels in MDA-EV and MDA-Tpm2.1 cells under both attached and suspension conditions. EphA2 expression decreased in suspension relative to attached conditions in both cell types and was further reduced in MDA-Tpm2.1 cells compared to MDA-EV cells under suspension conditions (Fig. 5F, G).

EphA2 can also be activated via a non-canonical mechanism involving AKT-mediated phosphorylation at serine 897 (Ser897). Western blot analysis revealed reduced levels of phospho-EphA2 (Ser897) in MDA-Tpm2.1 cells relative to MDA-EV cells under both attached and suspension conditions (Fig. S2). Supporting a pro-survival role for EphA2 signaling, treatment with an EphA2 kinase inhibitor significantly increased apoptosis in MDA-EV cells in suspension, to levels comparable to those observed in MDA-Tpm2.1 cells (Fig. 5H).

Together, these results suggest that AKT and EphA2 signaling promote anoikis resistance in breast cancer cells under anchorage-independent conditions and that suppression of these pathways by Tpm2.1 enhances apoptotic sensitivity in suspension.

## Discussion

Resistance to anoikis is a hallmark of metastatic cancer cells, allowing them to survive outside their native tissue environment^12^. While previous studies have demonstrated the role of Tpm2.1 in upregulating intrinsic apoptotic proteins^70^, this study reveals that restoring rigidity sensing in MDA-MB-231 cells through Tpm2.1 overexpression unmasks divergent transcriptional responses under suspension conditions. These responses define distinct fates, with some cells undergoing apoptosis while others survive by activating pro-survival mechanisms.

Our findings reveal that rigidity sensing alone is insufficient to guarantee apoptotic commitment in matrix-deprived breast cancer cells. While Tpm2.1 re-expression disrupts canonical survival signaling, only a fraction of cells fully undergo anoikis. This divergence reflects differential engagement of compensatory pathways. Cells with restored rigidity sensing can survive in suspension by activating a coordinated response involving immune signaling and intercellular adhesion. Together, these features suggest a shift toward cytokine-supported collective survival, enabling resilience under anchorage-independent conditions.

Conversely, cells that succumb to anoikis display a failure to activate these compensatory circuits. Instead of restoring pro-survival pathways, they remain transcriptionally repressed in critical regulators of adhesion and replication. Notably, their late upregulation of innate immune signaling may represent a stress-associated response that reinforces rather than prevents apoptosis. These contrasting trajectories underscore how cell fate under detachment stress hinges not only on the presence of rigidity cues but also on the cell’s capacity to restructure its transcriptional program toward survival. A schematic working model (Fig. 6) illustrates the distinct signaling pathways activated in each subpopulation of suspension-grown cells and their corresponding fates.

**Figure 6.**
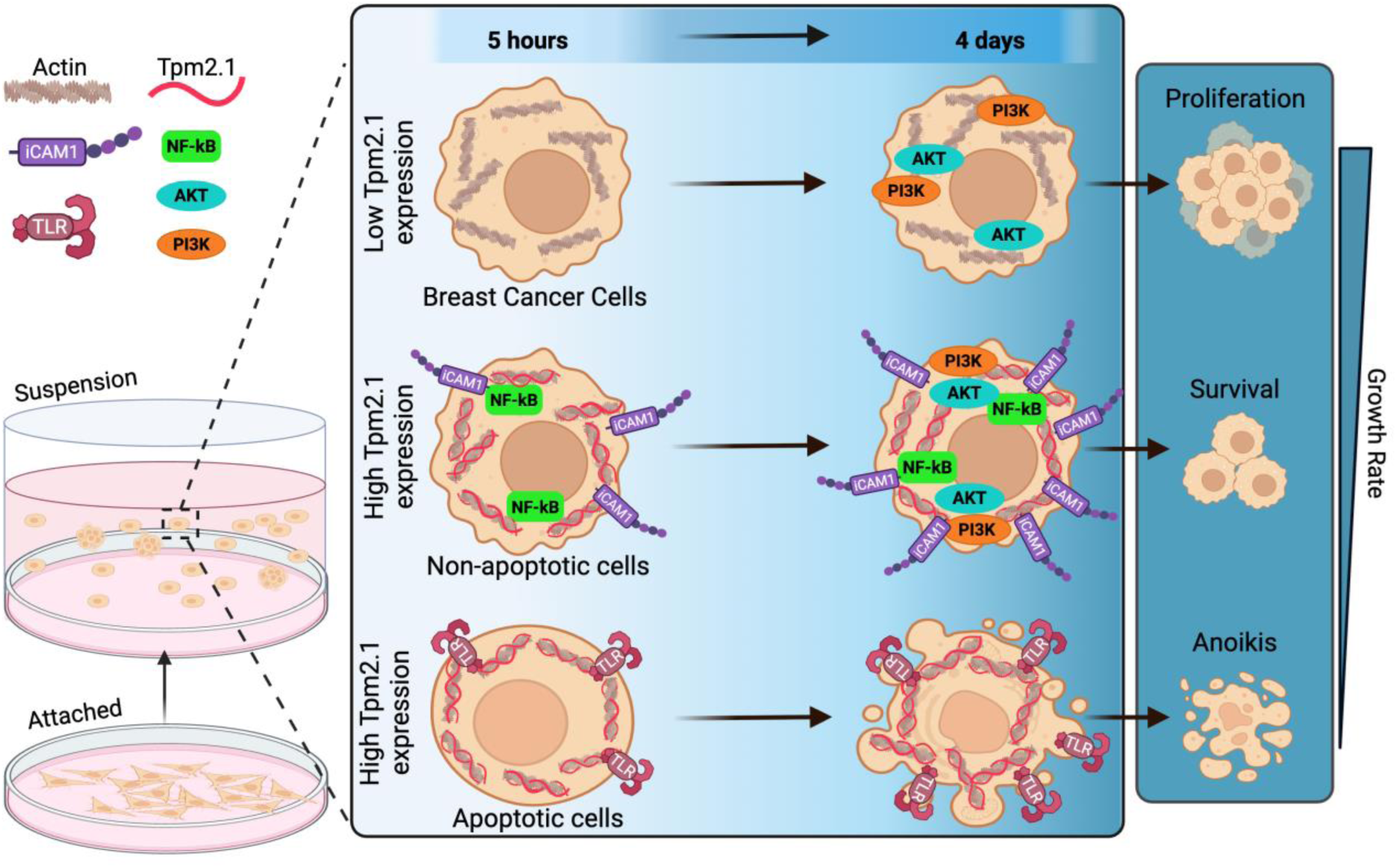
Illustrated Working Model. Schematic model depicting the distinct signaling pathways activated in breast cancer cells (MDA-EV) and Tpm2.1-expressing cells (MDA-Tpm2.1), following their separation into non-apoptotic (TpmNoAp) and apoptotic (TpmApop) subpopulations after 5 hours and 4 days in suspension. The model highlights subpopulation-specific pathway adaptations and the associated cellular fates under anchorage-independent conditions. Created in https://BioRender.com.

Mechanistically, we show that Tpm2.1 suppresses AKT phosphorylation in suspension without altering total AKT levels. Pharmacological inhibition of AKT increased apoptosis in both MDA-EV and Tpm2.1-expressing cells, confirming its role in promoting survival under anchorage-independent conditions. MDA-EV cells were especially sensitive to early AKT inhibition, while Tpm2.1-expressing cells were uniformly sensitive regardless of timing. These findings indicate that AKT is a key regulator of anoikis resistance and that its inhibition may be particularly effective during early suspension stress.

We also implicate EphA2, a known downstream effector of AKT^71^, in the regulation of cell survival. EphA2 expression and phosphorylation at Serine 897 residue were significantly reduced in Tpm2.1-overexpressing cells. Inhibition of EphA2 kinase activity phenocopied the pro-apoptotic effects of AKT suppression, suggesting that this receptor contributes to anchorage-independent survival in part through AKT-dependent signaling.

A further notable observation is the enhanced multicellular clustering of Tpm2.1-expressing cells in suspension. This morphological difference aligns with transcriptomic findings showing upregulation of adhesion-related genes in the non-apoptotic population. Elevated expression of ICAM1, in particular, has been associated with increased cell cohesion and protection from detachment-induced apoptosis. These findings suggest that in the presence of restored rigidity sensing, some cancer cells may adapt by reinforcing intercellular adhesion to maintain survival.

Together, this study provides a comprehensive view of how rigidity sensing and survival signaling intersect during detachment stress. While Tpm2.1 re-expression sensitizes cells to anoikis by disrupting pro-survival AKT and EphA2 pathways, a subset of cells compensate by activating inflammatory and adhesion networks. These findings highlight a dual mechanism of resistance involving both signaling rewiring and physical cell clustering. Targeting these compensatory circuits may provide new strategies to eliminate metastatic cancer cells that persist under matrix-deprived conditions.

## Materials and Methods

### Cell culture

MDA-MB-231 cells (wild-type, control, and Tpm2.1 overexpressed) were a generous gift from Dr. H. Wolfenson (Technion, Israel) and were originally obtained from ATCC^35^. Tpm2.1-overexpressing cells (MDA-Tpm) were generated by lentiviral overexpression of Tpm2.1 in MDA-MB-231 cells. All Cells were maintained in DMEM supplemented with 10% (v/v) fetal bovine serum (FBS), 1% penicillin–streptomycin, and 2 µg/mL puromycin at 37 °C in a humidified incubator with 5% CO₂.

To assess growth under non-adherent conditions, cells were cultured in ultra-low attachment plates or flasks (Corning, Cat. No. 3471 and 3814). Growth under suspension conditions serves as an alternative assay for cellular transformation and correlates strongly with results from soft agar assays^42^. For adherent conditions, cells were cultured on standard tissue culture-treated plates or flasks.

### Separation of apoptotic and non-apoptotic cells

MDA-Tpm2.1 cells were cultured under suspension conditions in ultra-low attachment plates for varying time points (5 h, 1 day, 2 days, and 4 days). Apoptotic and non-apoptotic populations were separated using the Annexin V MicroBead Kit (Miltenyi Biotec) according to the manufacturer’s protocol, which isolates cells based on Annexin V binding, a marker of early apoptosis^46^.

### Apoptosis measurement

Cells were cultured under adherent or suspension conditions for the indicated time periods. Adherent cells were harvested by trypsinization. All cells were collected by centrifugation, washed with DPBS, and resuspended in 100 µL Annexin V binding buffer. Apoptosis was assessed using Alexa Fluor™ 647-conjugated Annexin V (BD Pharmingen, Cat. No. 567356) and propidium iodide (PI; Invitrogen, Cat. No. BMS500F1-300), following the manufacturers’ protocols. Annexin V- and PI-positive populations were quantified by flow cytometry (BD FACSymphony A5) and analyzed using FlowJo software.

### Cell growth assay

Cells (5 × 10⁴ cells per well) were seeded in 6-well ultra-low attachment plates and cultured for 5 or 8 days. Cell viability was assessed using Trypan Blue exclusion, and live/dead cell counts were obtained using a DeNovix CellDrop FL cell counter. Fold change in cell number was calculated by dividing the number of live cells on Day 5 or Day 8 by the initial seeding density.

### Transwell-Migration Assay

MDA-EV and MDA-Tpm2.1 cells (1 × 10⁵ per well) were seeded in serum-free medium into the upper chamber of Transwell inserts (8 μm pore size; Corning Costar, catalog no. 3464) and incubated for 48 h at 37 °C in a humidified incubator with 5% CO₂. The lower chamber contained 500 μl of medium supplemented with 10% fetal bovine serum (FBS) as a chemoattractant. After incubation, the inserts were removed, and cells that had migrated to the lower surface of the membrane were fixed with 4% paraformaldehyde and stained with Hoechst dye. Phase-contrast and DAPI fluorescence images were acquired, and migrated cells were quantified using ImageJ (NIH).

### Inhibitor treatments

Cells were treated with the following small-molecule inhibitors: an AKT kinase inhibitor (Afuresertib Hydrochloride, 5 µM; MedChemExpress, Cat. No. HY-15727A/CS-3385), an ICAM-1 inhibitor (ICAM-1-IN-1, 100 nM; MedChemExpress, Cat. No. HY-U00003), and EphA2 kinase inhibitor (ALW-II-41-27, 1.5nM; MedChemExpress, Cat. No. HY18007), as indicated in each experiment.

### Western Blot

Cells were cultured in suspension (ultra-low attachment plates) or adherent (tissue culture plates) conditions for 5 days. Total protein was extracted using RIPA buffer (Cell Signaling Technology, Cat. No. 9806S) supplemented with protease inhibitors (Sigma-Aldrich, Cat. No. 11697498001) and incubated on ice for 45 minutes. Lysates were mixed with Laemmli buffer, separated on 4– 20% gradient SDS–PAGE gels (Bio-Rad, Cat. No. 4561096) at 120 V for 90 minutes, and transferred to nitrocellulose membranes using the Trans-Blot Turbo Transfer System (Bio-Rad).

Membranes were blocked in 5% BSA in 1× TBST for 1 hour at room temperature, followed by overnight incubation at 4 °C with primary antibodies. After washing, membranes were incubated with HRP-conjugated secondary antibodies for 1 hour at room temperature. The following primary antibodies were used: Anti-Tropomyosin (clone TM311, Sigma-Aldrich, T2780; 1:1000), AKT (pan) (Cell Signaling Technology, C67E7; 1:1000), phospho-AKT (T308) (Cell Signaling Technology, 244F9; 1:500), EphA2 (D4A2) (Cell Signaling Technology, 6997S; 1:1000), phoshpho-EphA2(S897) (Cell Signaling Technology, 6347S; 1:1000) and Cyclophilin B (Cell Signaling Technology, 43603S; 1:1000), used as a loading control. Secondary antibodies included HRP-conjugated anti-mouse IgG (EMD Millipore, AP160P) and anti-rabbit IgG (Santa Cruz Biotechnology, sc-2357).

### RNA isolation

Total RNA was isolated using the RNeasy Mini Kit (Qiagen®, Germantown, MD, USA; Cat. #74104) according to the manufacturer’s protocol. RNA concentration and purity were assessed using a NanoDrop Spectrophotometer (Thermo Fisher Scientific). A total of 28 RNA samples, representing 14 experimental conditions with biological duplicates, were submitted to Novogene Co. Ltd. (Nanjing, China) for library preparation and high-throughput sequencing.

### RNA sequencing and analysis

#### 1. Library preparation for Transcriptome sequencing- Non strand specific library

Messenger RNA was purified from total RNA using poly-T oligo-attached magnetic beads. After fragmentation, the first strand cDNA was synthesized using random hexamer primers followed by the second strand cDNA synthesis. The library was ready after end repair, A-tailing, adapter ligation, size selection, amplification, and purification. The library was checked with Qubit and real-time PCR for quantification and bioanalyzer for size distribution detection.

#### 2. Clustering and sequencing

After library quality control, different libraries were pooled based on the effective concentration and targeted data amount, then subjected to Illumina sequencing on the NovaSeq6000 Platform, using paired-end 150bp read length. The basic principle of sequencing is "Sequencing by Synthesis", where fluorescently labeled dNTPs, DNA polymerase, and adapter primers are added to the sequencing flow cell for amplification. As each sequencing cluster extends its complementary strand, the addition of each fluorescently labeled dNTP releases a corresponding fluorescence signal. The sequencer captures these fluorescence signals and converts them into sequencing peaks through computer software, thereby obtaining the sequence information of the target fragment.

#### 3. RNA Analysis

Differential expression analysis was performed using Novogene’s Novomagic platform, employing the DESeq2 pipeline with adjusted p-value (padj) < 0.05 and |log₂FoldChange| > 1 as significance thresholds.

Heatmap produced by an R script, with padj<0.05 and |log2Fold|>1, applying Spearman correlation and the centroid method for clustering.

## Acknowledgments

The authors would like to thank Dr. H. Wolfenson (Technion, Israel), Dr. Elisabeth Nadjar-Boger, Lidan Shi, and Malak Amer for kindly providing the MDA-MB-231 cell lines and for their helpful discussions. We are also grateful to Emma Pfortmiller and Dr. Michelle Ward (UTMB, USA) for their assistance in generating heatmaps and for their valuable input. Special thanks to Meredith Weglarz and the UTMB Flow Cytometry & Cell Sorting Core for their support with cytometry measurements and insightful discussions. This research was supported by UTMB startup funds and the Cancer Prevention & Research Institute of Texas (CPRIT) grant ID RR210018 to GN, a CPRIT scholar. Additional support was provided through postdoctoral fellowships from the Council for Higher Education of Israel and Bar-Ilan University, Israel, to AGV.

## References

1 V. H. Freedman, S.-i. Shin, Cellular tumorigenicity in nude mice: Correlation with cell growth in semi-solid medium. Cell 3, 355–359 (1974).

2 M. Sheetz, A Tale of Two States: Normal and Transformed, With and Without Rigidity Sensing. Annu Rev Cell Dev Biol 35, 169–190 (2019).

3 D. R. Welch, D. R. Hurst, Defining the Hallmarks of Metastasis. Cancer Res 79, 3011–3027 (2019).

4 J. Fares, M. Y. Fares, H. H. Khachfe, H. A. Salhab, Y. Fares, Molecular principles of metastasis: a hallmark of cancer revisited. Signal Transduction and Targeted Therapy 5, 28 (2020).

5 D. E. Discher, P. Janmey, Y.-l. Wang, Tissue Cells Feel and Respond to the Stiffness of Their Substrate. Science 310, 1139–1143 (2005).

6 B. Yang, H. Wolfenson, V. Y. Chung, N. Nakazawa, S. Liu, J. Hu, R. Y. Huang, M. P. Sheetz, Stopping transformed cancer cell growth by rigidity sensing. Nat Mater 19, 239–250 (2020).

7 A. Saraswathibhatla, D. Indana, O. Chaudhuri, Cell-extracellular matrix mechanotransduction in 3D. Nat Rev Mol Cell Biol 24, 495–516 (2023).

8 J. E. Meredith, B. Fazeli, M. A. Schwartz, The extracellular matrix as a cell survival factor. Molecular Biology of the Cell 4, 953–961 (1993).

9 S. M. Frisch, H. Francis, Disruption of epithelial cell-matrix interactions induces apoptosis. J Cell Biol 124, 619–626 (1994).

10 M. Taddei, E. Giannoni, T. Fiaschi, P. Chiarugi, Anoikis: an emerging hallmark in health and diseases. The Journal of Pathology 226, 380–393 (2012).

11 H. Wolfenson, B. Yang, M. P. Sheetz, Steps in Mechanotransduction Pathways that Control Cell Morphology. Annu Rev Physiol 81, 585–605 (2019).

12 Y. Dai, X. Zhang, Y. Ou, L. Zou, D. Zhang, Q. Yang, Y. Qin, X. Du, W. Li, Z. Yuan, Z. Xiao, Q. Wen, Anoikis resistance--protagonists of breast cancer cells survive and metastasize after ECM detachment. Cell Commun Signal 21, 190 (2023).

13 C. G. Galbraith, K. M. Yamada, M. P. Sheetz, The relationship between force and focal complex development. J Cell Biol 159, 695–705 (2002).

14 Sergey V. Plotnikov, Ana M. Pasapera, B. Sabass, Clare M. Waterman, Force Fluctuations within Focal Adhesions Mediate ECM-Rigidity Sensing to Guide Directed Cell Migration. Cell 151, 1513–1527 (2012).

15 A. Elosegui-Artola, E. Bazellières, M. D. Allen, I. Andreu, R. Oria, R. Sunyer, J. J. Gomm, J. F. Marshall, J. L. Jones, X. Trepat, P. Roca-Cusachs, Rigidity sensing and adaptation through regulation of integrin types. Nature Materials 13, 631–637 (2014).

16 T. Iskratsch, H. Wolfenson, M. P. Sheetz, Appreciating force and shape—the rise of mechanotransduction in cell biology. Nat Rev Mol Cell Biol 15, 825–833 (2014).

17 V. Vogel, M. P. Sheetz, Cell fate regulation by coupling mechanical cycles to biochemical signaling pathways. Current Opinion in Cell Biology 21, 38–46 (2009).

18 Y. H. Zhang, C. Q. Zhao, L. S. Jiang, L. Y. Dai, Substrate stiffness regulates apoptosis and the mRNA expression of extracellular matrix regulatory genes in the rat annular cells. Matrix Biol 30, 135–144 (2011).

19 G. Meacci, H. Wolfenson, S. Liu, M. R. Stachowiak, T. Iskratsch, A. Mathur, S. Ghassemi, N. Gauthier, E. Tabdanov, J. Lohner, A. Gondarenko, A. C. Chander, P. Roca-Cusachs, B. O’Shaughnessy, J. Hone, M. P. Sheetz, α-Actinin links extracellular matrix rigidity-sensing contractile units with periodic cell-edge retractions. Mol Biol Cell 27, 3471–3479 (2016).

20 A. Kumar, J. K. Placone, A. J. Engler, Understanding the extracellular forces that determine cell fate and maintenance. Development 144, 4261–4270 (2017).

21 D. W. Zhou, T. T. Lee, S. Weng, J. Fu, A. J. García, Effects of substrate stiffness and actomyosin contractility on coupling between force transmission and vinculin-paxillin recruitment at single focal adhesions. Mol Biol Cell 28, 1901–1911 (2017).

22 H. Shin, D. Kim, D. M. Helfman, Tropomyosin isoform Tpm2.1 regulates collective and amoeboid cell migration and cell aggregation in breast epithelial cells. Oncotarget 8, 95192–95205 (2017).

23 X. Zhou, Z. Li, H. Chen, M. Jiao, C. Zhou, H. Li, Relevance Analysis of TPM2 and Clinicopathological Characteristics in Breast Cancer. Int J Gen Med 17, 59–74 (2024).

24 H. Wolfenson, G. Meacci, S. Liu, M. R. Stachowiak, T. Iskratsch, S. Ghassemi, P. Roca-Cusachs, B. O’Shaughnessy, J. Hone, M. P. Sheetz, Tropomyosin controls sarcomere-like contractions for rigidity sensing and suppressing growth on soft matrices. Nat Cell Biol 18, 33–42 (2016).

25 Q. Chen, M. S. Kinch, T. H. Lin, K. Burridge, R. L. Juliano, Integrin-mediated cell adhesion activates mitogen-activated protein kinases. J Biol Chem 269, 26602–26605 (1994).

26 D. D. Schlaepfer, S. K. Hanks, T. Hunter, P. van der Geer, Integrin-mediated signal transduction linked to Ras pathway by GRB2 binding to focal adhesion kinase. Nature 372, 786–791 (1994).

27 X. Pang, X. He, Z. Qiu, H. Zhang, R. Xie, Z. Liu, Y. Gu, N. Zhao, Q. Xiang, Y. Cui, Targeting integrin pathways: mechanisms and advances in therapy. Signal Transduction and Targeted Therapy 8, 1 (2023).

28 G. S. Goldberg, Z. Jin, H. Ichikawa, A. Naito, M. Ohki, W. S. El-Deiry, H. Tsuda, Global Effects of Anchorage on Gene Expression during Mammary Carcinoma Cell Growth Reveal Role of Tumor Necrosis Factor-related Apoptosis-inducing Ligand in Anoikis1. Cancer Research 61, 1334–1337 (2001).

29 O. W. Merten, Advances in cell culture: anchorage dependence. Philos Trans R Soc Lond B Biol Sci 370, 20140040 (2015).

30 H. D. Huh, Y. Sub, J. Oh, Y. E. Kim, J. Y. Lee, H.-R. Kim, S. Lee, H. Lee, S. Pak, S. E. Amos, D. Vahala, J. H. Park, J. E. Shin, S. Y. Park, H. S. Kim, Y. H. Roh, H.-W. Lee, K.-L. Guan, Y. S. Choi, J. Jeong, J. Choi, J.-S. Roe, H. Y. Gee, H. W. Park, Reprogramming anchorage dependency by adherent-to-suspension transition promotes metastatic dissemination. Molecular Cancer 22, 63 (2023).

31 F. O. Adeshakin, A. O. Adeshakin, L. O. Afolabi, D. Yan, G. Zhang, X. Wan, Mechanisms for Modulating Anoikis Resistance in Cancer and the Relevance of Metabolic Reprogramming. Front Oncol 11, 626577 (2021).

32 D. Fanfone, Z. Wu, J. Mammi, K. Berthenet, D. Neves, K. Weber, A. Halaburkova, F. Virard, F. Bunel, C. Jamard, H. Hernandez-Vargas, S. W. G. Tait, A. Hennino, G. Ichim, Confined migration promotes cancer metastasis through resistance to anoikis and increased invasiveness. Elife 11 (2022).

33 G. N. Raval, S. Bharadwaj, E. A. Levine, M. C. Willingham, R. L. Geary, T. Kute, G. L. Prasad, Loss of expression of tropomyosin-1, a novel class II tumor suppressor that induces anoikis, in primary breast tumors. Oncogene 22, 6194–6203 (2003).

34 R. Qin, S. Melamed, B. Yang, M. Saxena, M. P. Sheetz, H. Wolfenson, Tumor Suppressor DAPK1 Catalyzes Adhesion Assembly on Rigid but Anoikis on Soft Matrices. Frontiers in Cell and Developmental Biology 10 (2022).

35 L. Shi, E. Nadjar-Boger, H. Jafarinia, A. Carlier, H. Wolfenson, YAP mediates apoptosis through failed integrin adhesion reinforcement. Cell reports 43 (2024).

36 T. Liu, L. Zhang, D. Joo, S.-C. Sun, NF-κB signaling in inflammation. Signal Transduction and Targeted Therapy 2, 17023 (2017).

37 O. D. Perez, S. Kinoshita, Y. Hitoshi, D. G. Payan, T. Kitamura, G. P. Nolan, J. B. Lorens, Activation of the PKB/AKT pathway by ICAM-2. Immunity 16, 51–65 (2002).

38 R. Liu, Y. Chen, G. Liu, C. Li, Y. Song, Z. Cao, W. Li, J. Hu, C. Lu, Y. Liu, PI3K/AKT pathway as a key link modulates the multidrug resistance of cancers. Cell Death & Disease 11, 797 (2020).

39 Y. He, M. M. Sun, G. G. Zhang, J. Yang, K. S. Chen, W. W. Xu, B. Li, Targeting PI3K/Akt signal transduction for cancer therapy. Signal Transduction and Targeted Therapy 6, 425 (2021).

40 X. Hou, C. Ren, J. Jin, Y. Chen, X. Lyu, K. Bi, N. D. Carrillo, V. L. Cryns, R. A. Anderson, J. Sun, M. Chen, Phosphoinositide signalling in cell motility and adhesion. Nature Cell Biology 27, 736–748 (2025).

41 B. Izar, A. Rotem, GILA, a Replacement for the Soft-Agar Assay that Permits High-Throughput Drug and Genetic Screens for Cellular Transformation. Curr Protoc Mol Biol 116, 28.28.21–28.28.12 (2016).

42 A. Rotem, A. Janzer, B. Izar, Z. Ji, J. G. Doench, L. A. Garraway, K. Struhl, Alternative to the soft-agar assay that permits high-throughput drug and genetic screens for cellular transformation. Proceedings of the National Academy of Sciences 112, 5708–5713 (2015).

43 J. Yang, S. A. Mani, J. L. Donaher, S. Ramaswamy, R. A. Itzykson, C. Come, P. Savagner, I. Gitelman, A. Richardson, R. A. Weinberg, Twist, a Master Regulator of Morphogenesis, Plays an Essential Role in Tumor Metastasis. Cell 117, 927–939 (2004).

44 A. J. Minn, G. P. Gupta, P. M. Siegel, P. D. Bos, W. Shu, D. D. Giri, A. Viale, A. B. Olshen, W. L. Gerald, J. Massagué, Genes that mediate breast cancer metastasis to lung. Nature 436, 518–524 (2005).

45 O. Friedrich, D. F. Gilbert, Cell viability assays : methods and protocols, Methods in molecular biology, 2644 (Humana Press, New York, NY, ed. Second edition., 2023), 10.1007/978-1-0716-3052-5.

46 A.-M. Lobascio, F.-G. Klinger, M. De Felici, Isolation of apoptotic mouse fetal oocytes by AnnexinV assay. International Journal of Developmental Biology 51 (2007).

47 M. Kanehisa (2002) The KEGG database. in ‘In silico’simulation of biological processes: Novartis Foundation Symposium 247 (Wiley Online Library), pp 91–103.

48 C. J. Antalis, A. Uchida, K. K. Buhman, R. A. Siddiqui, Migration of MDA-MB-231 breast cancer cells depends on the availability of exogenous lipids and cholesterol esterification. Clin Exp Metastasis 28, 733–741 (2011).

49 C. Rodrigues dos Santos, G. Domingues, I. Matias, J. Matos, I. Fonseca, J. M. de Almeida, S. Dias, LDL-cholesterol signaling induces breast cancer proliferation and invasion. Lipids in Health and Disease 13, 16 (2014).

50 T. Aasen, M. Mesnil, C. C. Naus, P. D. Lampe, D. W. Laird, Gap junctions and cancer: communicating for 50 years. Nat Rev Cancer 16, 775–788 (2016).

51 D. Kyuno, A. Takasawa, S. Kikuchi, I. Takemasa, M. Osanai, T. Kojima, Role of tight junctions in the epithelial-to-mesenchymal transition of cancer cells. Biochimica et Biophysica Acta (BBA) - Biomembranes 1863, 183503 (2021).

52 I. Baroja, N. C. Kyriakidis, G. Halder, I. M. Moya, Expected and unexpected effects after systemic inhibition of Hippo transcriptional output in cancer. Nature Communications 15, 2700 (2024).

53 R. Jinka, R. Kapoor, P. G. Sistla, T. A. Raj, G. Pande, Alterations in Cell-Extracellular Matrix Interactions during Progression of Cancers. Int J Cell Biol 2012, 219196 (2012).

54 L. H. Xu, X. Yang, C. A. Bradham, D. A. Brenner, A. S. Baldwin, Jr., R. J. Craven, W. G. Cance, The focal adhesion kinase suppresses transformation-associated, anchorage-independent apoptosis in human breast cancer cells. Involvement of death receptor-related signaling pathways. J Biol Chem 275, 30597–30604 (2000).

55 H. Akca, A. Demiray, O. Tokgun, J. Yokota, Invasiveness and anchorage independent growth ability augmented by PTEN inactivation through the PI3K/AKT/NFkB pathway in lung cancer cells. Lung Cancer 73, 302–309 (2011).

56 K. Tsirtsaki, V. Gkretsi, The focal adhesion protein Integrin-Linked Kinase (ILK) as an important player in breast cancer pathogenesis. Cell Adh Migr 14, 204–213 (2020).

57 P. Paoli, E. Giannoni, P. Chiarugi, Anoikis molecular pathways and its role in cancer progression. Biochimica et Biophysica Acta (BBA) - Molecular Cell Research 1833, 3481–3498 (2013).

58 W. Zhang, H. T. Liu, MAPK signal pathways in the regulation of cell proliferation in mammalian cells. Cell Research 12, 9–18 (2002).

59 J. Yue, J. M. López, Understanding MAPK Signaling Pathways in Apoptosis. Int J Mol Sci 21 (2020).

60 X. Zhong, F. J. Rescorla, Cell surface adhesion molecules and adhesion-initiated signaling: Understanding of anoikis resistance mechanisms and therapeutic opportunities. Cellular Signalling 24, 393–401 (2012).

61 H. Zhang, L. Zhang, M. Lu, Inhibition of integrin subunit alpha 11 restrains gastric cancer progression through phosphatidylinositol 3-kinase/Akt pathway. Bioengineered 12, 11909–11921 (2021).

62 G. Huang, M. Zhou, D. Lu, J. Li, Q. Tang, C. Xiong, F. Liang, R. Chen, The mechanism of ITGB4 in tumor migration and invasion. Frontiers in Oncology 14 (2024).

63 H. Li, H. Liu, L. Xiao, H. Gao, H. Wei, A. Han, G. Lin, A Novel Oncogenic and Drug-Sensitive KIF5B-NTRK1 Fusion in Lung Adenocarcinoma. Curr Oncol 31, 6621–6631 (2024).

64 A. O. Aliprantis, R. B. Yang, D. S. Weiss, P. Godowski, A. Zychlinsky, The apoptotic signaling pathway activated by Toll-like receptor-2. Embo j 19, 3325–3336 (2000).

65 H. Yi, A. K. Patel, C. P. Sodhi, D. J. Hackam, A. S. Hackam, Novel Role for the Innate Immune Receptor Toll-Like Receptor 4 (TLR4) in the Regulation of the Wnt Signaling Pathway and Photoreceptor Apoptosis. PLOS ONE 7, e36560 (2012).

66 K. Czubak-Prowizor, A. Babinska, M. Swiatkowska, The F11 Receptor (F11R)/Junctional Adhesion Molecule-A (JAM-A) (F11R/JAM-A) in cancer progression. Mol Cell Biochem 477, 79–98 (2022).

67 C. Guerra-Espinosa, M. Jiménez-Fernández, F. Sánchez-Madrid, J. M. Serrador, ICAMs in Immunity, Intercellular Adhesion and Communication. Cells 13 (2024).

68 R. Taftaf, X. Liu, S. Singh, Y. Jia, N. K. Dashzeveg, A. D. Hoffmann, L. El-Shennawy, E. K. Ramos, V. Adorno-Cruz, E. J. Schuster, D. Scholten, D. Patel, Y. Zhang, A. A. Davis, C. Reduzzi, Y. Cao, P. D’Amico, Y. Shen, M. Cristofanilli, W. A. Muller, V. Varadan, H. Liu, ICAM1 initiates CTC cluster formation and trans-endothelial migration in lung metastasis of breast cancer. Nature Communications 12, 4867 (2021).

69 B. D. Manning, A. Toker, AKT/PKB Signaling: Navigating the Network. Cell 169, 381–405 (2017).

70 M. Desouza-Armstrong, P. W. Gunning, J. R. Stehn, Tumor suppressor tropomyosin Tpm2.1 regulates sensitivity to apoptosis beyond anoikis characterized by changes in the levels of intrinsic apoptosis proteins. Cytoskeleton 74, 233–248 (2017).

71 H. Miao, D. Q. Li, A. Mukherjee, H. Guo, A. Petty, J. Cutter, J. P. Basilion, J. Sedor, J. Wu, D. Danielpour, A. E. Sloan, M. L. Cohen, B. Wang, EphA2 mediates ligand-dependent inhibition and ligand-independent promotion of cell migration and invasion via a reciprocal regulatory loop with Akt. Cancer Cell 16, 9–20 (2009).

